# A Combined Numerical and Experimental Investigation of Localized Electroporation-based Transfection and Sampling

**DOI:** 10.1101/346981

**Authors:** Prithvijit Mukherjee, S. Shiva P. Nathamgari, John A. Kessler, Horacio D. Espinosa

## Abstract

Localized electroporation has evolved as an effective technology for the delivery of foreign molecules into adherent cells, and more recently, for the sampling of cytosolic content from a small population of cells. Unlike bulk electroporation, where the electric field is poorly controlled, localized electroporation benefits from the spatial localization of the electric field on a small areal fraction of the cell membrane, resulting in efficient molecular transport and high cell-viability. Although there have been numerous experimental reports, a mechanistic understanding of the different parameters involved in localized electroporation is lacking. In this work, we developed a multiphysics model that a) predicts the electro-pore distribution in response to the local transmembrane potential and b) calculates the molecular transport into and out of the cell based on the predicted pore-sizes. Using the model, we identify that cell membrane tension plays a crucial role in enhancing both the amount and the uniformity of molecular transport, particularly for large proteins and plasmids. We qualitatively validate the model predictions by delivering large molecules (fluorescent-tagged bovine serum albumin and mCherry encoding plasmid) and by sampling an exogeneous protein (tdTomato) in an engineered cell line. The findings presented here should inform the future design of microfluidic devices for localized electroporation based sampling, eventually paving the way for temporal, single-cell analysis.

## INTRODUCTION

The ability to monitor internal cellular biomarkers at different time points and measure their changes over time is key to understanding the fundamental mechanisms governing dynamic cellular processes such as differentiation, maturation and ageing [1–3]. Temporal measurement of intracellular contents at the single-cell level can provide insights into the involved regulatory pathways and help understand the pathophysiology of disorders such as Alzheimer’s and Parkinson’s as well as the efficacy of drugs and their possible toxic effects [4–6].

Current high-throughput single-cell technologies for genomic, transcriptomic and proteomic analysis combine microfluidic platforms such as droplets, valves and nanowells [7] with high sensitivity assays such as single-cell western blot, protein barcodes, antibody spots and RNA sequencing [8–14]. These methods rely exclusively on cell lysis, and thus provide only a single snapshot of cellular activity in time. Using these lysis based methods, pseudo-time histories can be constructed with information obtained from parallel cultures [15]. The caveat to this approach is that cellular heterogeneity can mask the true temporal variations in biomarker levels. A commonly used technique of temporal investigation of the same cells involves the use of molecular probes (nanostars, molecular beacons and fluorescent biosensors) to tag molecules of interest in live cells [16–21]. However, the number of intracellular targets that can be studied simultaneously and the possibility of cellular perturbation caused by these intracellular labels limit the utility of these methods. Another approach for temporal analyses, which is limited to a sub-set of cellular proteins, uses microfluidic platforms to profile and record secreted proteins across a timespan [22, 23]. Recent approaches have also used nanoprobes such as carbon nanotubes, AFM tips and nanopipettes to extract single-cell cytosolic content for subsequent assays [24–27]. These methods however, are serial and lack the throughput required for systems biology analyses.

Micro and nano-scale electroporation technologies offer significant advantages over traditional bulk electroporation methods for the delivery of small molecules, proteins and nucleic acids of interest into cells [28–32]. The applied electric fields in these techniques are gentle to the cells and often perturb only a small fraction of the cell membrane. Two major advantages that these localized electroporation methods offer are the preservation of cell viability and functionality and the ability to target single-cells for delivery and subsequent monitoring [28, 33]. As such, electroporation at the micro and nano-scale has also been used to address the converse problem, i.e. to non-destructively sample cytosolic contents from small populations of cells [34, 35]. Although these techniques present promising approaches towards temporal sampling from live single-cells, there are major challenges that need to be overcome. One technological challenge that inhibits the realization of single-cell temporal sampling is the necessity of high precision microfluidic systems coupled to high sensitivity assays that can handle, transport and detect subcellular amounts of analytes in picoliter volumes, without incurring substantial losses. Another major hurdle is the lack of a mechanistic understanding of the process of localized electroporation and molecular transport out of the cell during sampling.

To improve our understanding of the process of localized electroporation and molecular transport, we have developed a multiphysics model incorporating the dynamics of pore formation on the cell membrane in response to a non-uniform and localized electric field; and the subsequent transport of molecules of interest into or out of the cells through these membrane pores. We have validated the model by quantifying the delivery and sampling of proteins in a small cell population using the so-called Localized Electroporation Device (LEPD) [29] – a microfluidic device developed by the Espinosa group for the culture and localized electroporation of adherent cells. The experimental trends corroborate with the model predictions and together they provide regimes of operation in the applied pulse strength and frequency, which are ideal for efficient delivery and sampling without compromising cell viability. The results also provide general guidelines regarding optimization of pulse parameters and device design applicable to localized electroporation mediated delivery and sampling. These guidelines lay down the foundations necessary to achieve the goal of single-cell temporal sampling.

## RESULTS AND DISCUSSIONS

### Device Architecture and Operation

The LEPD architecture allows for the long-term culture and localized electroporation of adherent cells. The cells are cultured on a polycarbonate substrate with multiple nanochannels that is sandwiched between a PDMS micro-well layer and a delivery/sampling chamber (**Figure 1a**). This chamber can serve the dual purpose of retaining the molecular cargo to be delivered into the cells or collecting the intracellular molecules that leak out from the cell during the process of electroporation. The extracted cytosolic content can then be retrieved for downstream analyses. The substrate material and nanochannel density can be varied according to experimental requirements. When an electric field is applied across the LEPD, the nanochannels in the substrate confine the electric field to a small fraction of the cell membrane and minimize perturbations to the cell state. Thus, this architecture can be used to transfect and culture sensitive cells (such as primary cells) while preserving a high degree of cell-viability. The Espinosa group has previously demonstrated on-chip differentiation of murine neural stem cells and transfection of postmitotic neurons on the LEPD platform [29]. In the current work, the LEPD has been extended to sampling an exogenous protein in a small population of engineered cells. All of the experimental data and computational analyses presented here were acquired using the LEPD architecture.

### Multiphysics Modeling

The application of pulsed electric fields leads to the transient permeabilization of the cell membrane, allowing both influx and outflux of molecules of varying sizes [36–38]. The most widely reported mechanism underlying this phenomenon assumes the formation of hydrophilic toroidal pores in the phospholipid bilayer [39, 40]. The formation, evolution and destruction of these ‘electro-pores’ is governed by the Smoluchowski advection-diffusion equation, which is derived using a statistical mechanics framework [41, 42]. Assuming the molecular transport through these membrane electro-pores to be primarily diffusive and electrophoretic, this framework has been utilized to provide estimates of small molecule delivery in bulk electroporation [43]. Although this model simplifies certain chemical and mechanical aspects of the permeabilization process [44–46], it correlates reasonably well with experimental observations of electropores [47–49], molecular dynamics simulations [50, 51] and molecular transport [43]. Here we extend this model to the case of localized electroporation where a spatially focused electric field is utilized to permeabilize only a small fraction of the cell membrane and deliver or extract molecules of interest. Specifically, we investigate the following mechanistic aspects of localized electroporation mediated transport: 1) The role of cell membrane tension in enhancing molecular transport; 2) The dependence of molecular transport (in both delivery and sampling) on the strength of the applied electric field; 3) The differences in delivery efficiency depending on molecular size. In the subsequent sections, we first discuss the implementation of the model followed by the results obtained.

#### Transmembrane Potential

Unlike bulk-electroporation where the electric field is maximum at the poles facing the electrodes and decays away from them[52–54], the electric field in localized electroporation is focused near the nanochannels and drops rapidly outside of them. By solving the electric field distribution (e.g. using the Finite Element Method), for the case of a cell placed on a substrate with a single nanochannel (radius = 250 nm) underneath, we found that the TMP drops rapidly (to 1/*e* of the maximum value within ~1.5 times the radius of the nanochannel) outside the region where the cell membrane interfaces the nanochannel (**see Figure 1c**). Since the electric field is confined to the region of the nanochannel, the concept can be extended to multiple nanochannels and an equivalent electrical circuit (**see Figure 1c**) can describe the LEPD system. The equivalent circuit model provides the flexibility of incorporating complex geometries, like the LEPD where several substrate nanochannels (~100 to 1000) can interface with a single cell, that are otherwise computationally non-trivial to simulate using a full field model. In the equivalent circuit *R_c_* and *C_c_* are the contact resistance and capacitance at the electrode-buffer interface. *R_s_* includes all the system resistances in series such as the buffer in the device and the external circuit. The top part of the cell membrane not interfacing with the nanochannels on the PC substrate is modeled by the resistive and capacitive elements *R_t_* and *C_t_*. The cell cytoplasm is represented by the resistance *R_cy_*. *R_b_* and *C_b_* represent the equivalent resistance and capacitance of the bottom cell membrane fraction interfacing the substrate nanochannels. *R_t_* and *R_b_* are variable resistors as the conductivity (κ) of the cell membrane changes with the formation and evolution of electro-pores (**see Supplementary S1**). Since the electric field is localized, only the bottom membrane fraction interfacing the nanochannels has been accounted for by these circuit elements. *R_g_* is the leakage resistance between the cells and the nanochannels. *R_cp_* and *R_op_* represent the equivalent resistance of all the nanochannels covered by the cells and the open nanochannels respectively. A system of three ODEs, derived using voltage and current conservation laws at each node, can be used to solve for the TMP, viz.
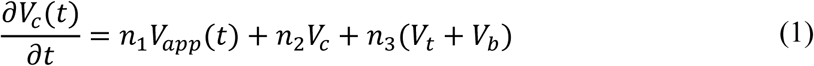

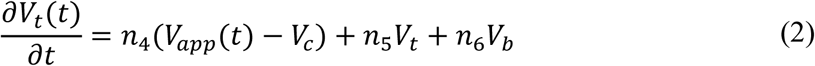

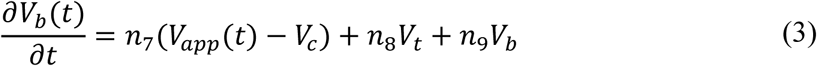

Here, *V_c_* is the potential drop across the contact, *V_t_* is the TMP across the top cell membrane, *V_b_* is the TMP across the bottom cell membrane and *V_m_* is the applied far-field voltage. Coefficients *n*_1_ to *n*_9_ are functions of the circuit elements (**see Supplementary Table 1**). The TMP values *V_t_* and *V_b_* are utilized in the pore evolution equation which is discussed next.

**Figure 1:**
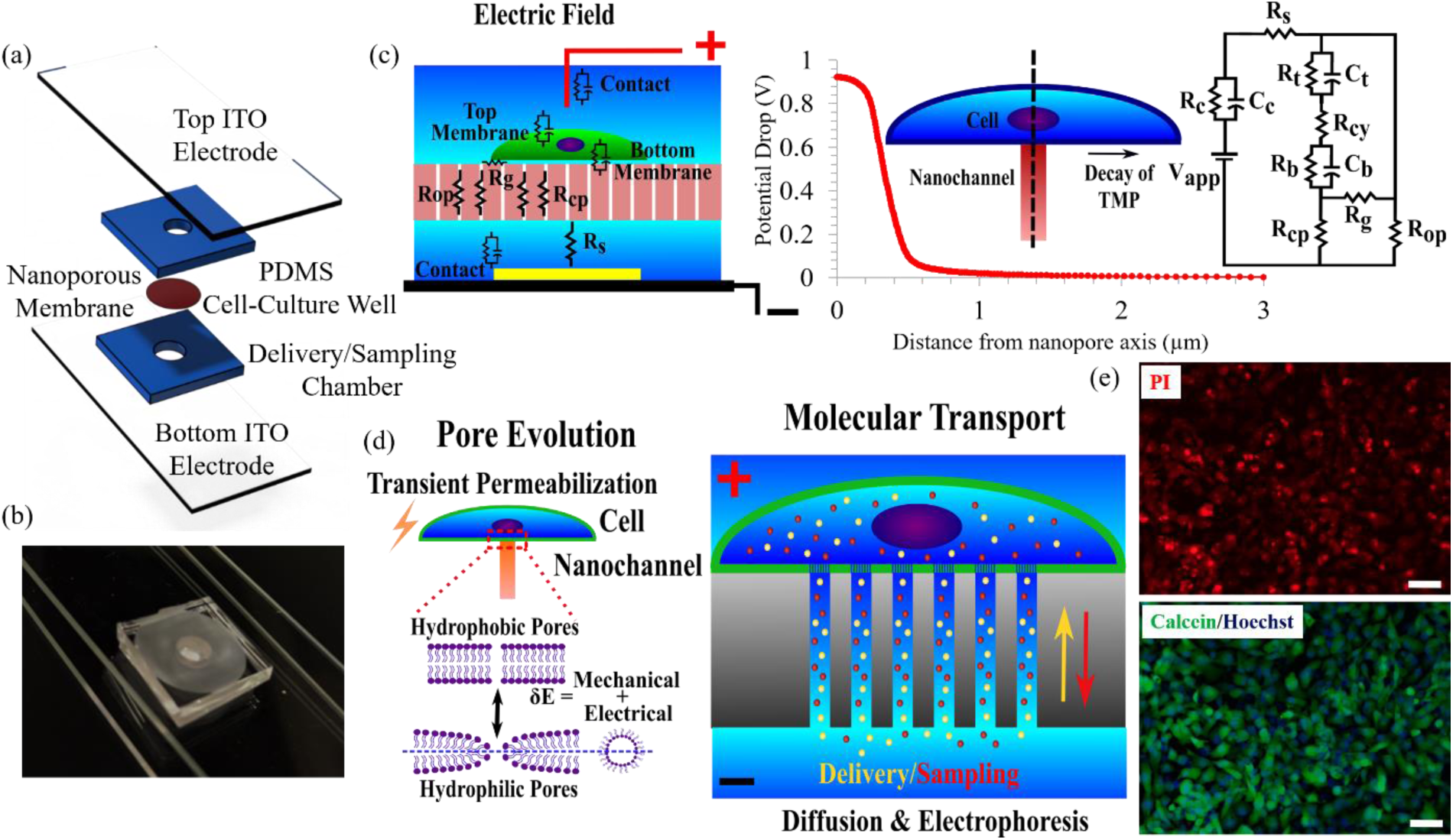
Overview of the experimental and computational framework (a) Schematic of the Localized Electroporation Device (LEPD) showing the different constituent layers, (b) Optical image of LEPD consisting of the PDMS device sandwiched between two ITO electrodes, (c) Left – Schematic of the concept of localized electroporation and the components that can be used to describe the electric field distribution. The transmembrane potential (TMP) is obtained by solving the electric field equations. Right – Axisymmetric FEM simulation of the electric field with a single nanochannel underneath a cell shows that the transmembrane potential drop is confined to the region of the nanochannel for localized electroporation. Consequently, a lumped circuit model can be used to represent a system with many nanochannels in parallel underneath a cell, (d) Left – Schematic of the pore evolution model. Transient permeabilization of the plasma membrane leads to the formation of hydrophilic pores that allow the passage of molecules. The size distribution of pores formed in response to an elevated TMP is obtained by solving a non-linear advection-diffusion equation. Right – Schematic of the molecular transport model. The transport across the permeabilized membrane is primarily diffusive and electrophoretic, (e) Delivery of PI into HT 1080 cells on the LEPD platform under iso-osmolar conditions using a 10 V pulse. Top image shows a delivery efficiency of > 95%. Bottom image shows viability of >95% using live dead staining, 6 hours post electroporation (Scale bars = 50 µm).

#### Pore Evolution

The evolution of electro-pores in bilayer membranes is governed by the Smoluchowski equation-
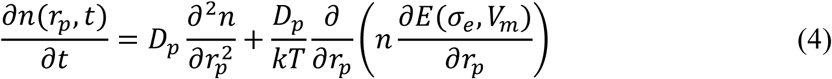

where, *n* is the density of electro-pores per unit membrane area between radius *r_p_* and *r_p_* + *dr_p_* at a particular time *t*. *D_p_* is the pore diffusion coefficient in the pore radius space *r_p_*, *kT* is the thermal energy and *E* is the energy difference between a bilayer membrane with and without a hydrophilic electro-pore (**see Supplementary S2**). The energy *E* is a function of the effective bilayer membrane tension *σ_e_* and the TMP *V_m_*, which in our system are the values *V_t_* and *V_b_* obtained by solving the equivalent electrical circuit. The effective membrane tension (*σ_e_*) assumes a non-linear form, which allows for the coupling of the electro-pores. The effective membrane tension (*σ_e_*) is a function of the surface tension (*σ*) of the cell membrane without pores and the pore distribution (*n*) (**Supplementary S2**). The pore evolution equation is solved in a one-dimensional radius space with appropriate pore creation and destruction rates (**Supplementary S3**) as the boundary condition [42] at the minimum pore radius (*r_min_*) and a no flux boundary condition at the maximum radius (*r_max_*), which represents the largest permissible electro-pore size. The pore evolution is coupled to the electric field by the effective membrane conductivity (κ) and the dynamics of electro-pores depend on the effective membrane tension (*σ_e_*). Briefly, the formation of electro-pores in response to the increased TMP (*V_t_*, *V_b_*) leads to an increase in the effective membrane conductivity (κ), since the electro-pores act as parallel ion-conducting pathways. The conductivity increase results in a drop of the cell membrane resistance represented by the circuit elements *R_t_* and *R_b_*. Consequently, the TMP decreases which arrests the nucleation and expansion of the electro-pores. The effective membrane tension (*σ_e_*) also drops as a result of the formed electro-pores [55, 56]. The reduction in membrane tension increases the energy required to expand the formed electro-pores, which eventually halts their growth. These interactions control the pore dynamics based on which we calculate the molecular transport across the permeabilized bottom cell membrane.

#### Molecular Transport

The molecular flux across the permeabilized cell membrane is calculated using the Nernst-Planck equation:
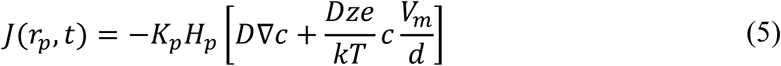

here, *J* is the flux of molecules across the membrane, *K_p_* and *H_p_* are the partition and hindrance factors[42, 43] respectively, *D* is the diffusion coefficient of the molecule of interest, *c* is the local concentration of the molecule, *z* is the charge on the molecule, *e* is the elementary charge, *kT* is the thermal energy, *V_m_* is the TMP and *d* is the thickness of the cell membrane. For our calculations, we have assumed that the concentration gradient does not change during the pulsation period. The total transported amount is calculated by integrating the flux through the electro-pores over time for the duration of the applied electric pulse, namely,
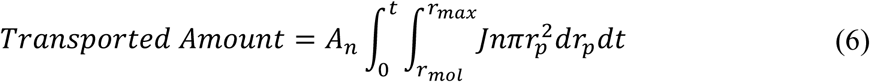

where, *A_n_* is the total cell membrane area interfacing with the nanochannels and *r_mol_* is the radius of the molecule of interest. For simplicity, it is assumed that only pores larger than the hydrodynamic radius of the molecule of interest permit their transport. The values of parameters used in the model are listed in the supplementary section (**Supplementary Table 2**).

### Model Predictions and Experimental Validation

#### Increasing the cell membrane tension enhances molecular transport

Previously, it has been shown that higher membrane tension facilitates electropermeabilization at lower electric field strengths in both lipid bilayers and mammalian cells that were bulk electroporated[57, 58]. The fact that membrane tension plays a critical role in cell membrane permeabilization is also evident from mechano-poration literature where a mechanically induced large deformation enables the permeabilization and subsequent transfection of cells[59, 60]. Naturally, we were interested in understanding the effect of modulating membrane tension in the case of localized electroporation.

We investigated the effect of membrane tension on the molecular transport of large molecules (hydrodynamic radius > 3 nm) in the LEPD system. Our analysis predicts that with an increase in the membrane tension, the transported amount for molecules larger than 3 nm in size increases for all applied voltages within a range (**see Figure 2b**). In our simulations, four different membrane tension values ranging from 1e-5 N/m to 8e-4 N/m were used. These values are within the range of membrane tensions reported in literature [61]. For each value of membrane tension, far-field voltages ranging from 7 V to 25 V were investigated. Below 7 V, electro-pores that enable the transport of molecules larger than 3 nm were not formed. It is important to note that the membrane tension referred to in this section is the initial tension for a cell membrane without pores (*σ*). The simulation results indicate two complementary mechanisms by which an increase in the membrane tension enhances molecular transport. First, the number of large pores per cell (>3 nm) as well as the mean radius of large pores (**see Figure 2c and Figure 2d**) increases at higher membrane tensions. In addition, a higher membrane tension stabilizes the large pores during the electric pulse application. This is confirmed by the existence of a greater number of large pores for longer duration during the applied electric pulse (**see Figure 2c**) and the quicker expansion of the pores to the largest radii (15 nm in our model) for high membrane tensions (**see Figure 2d**). Overall, the model results predict that increasing membrane tension can increase the efficiency of molecular transport even in the case of localized electroporation. This has indeed been observed in localized electroporation systems where suction pressure[62] or high aspect ratio nano-structures have been used to efficiently transfect cells[30]. We believe that in addition to reducing the electric field leakage due to imperfect sealing, these methods induce higher tension in the cell membrane, which facilitates efficient molecular delivery according to our analysis.

**Figure 2:**
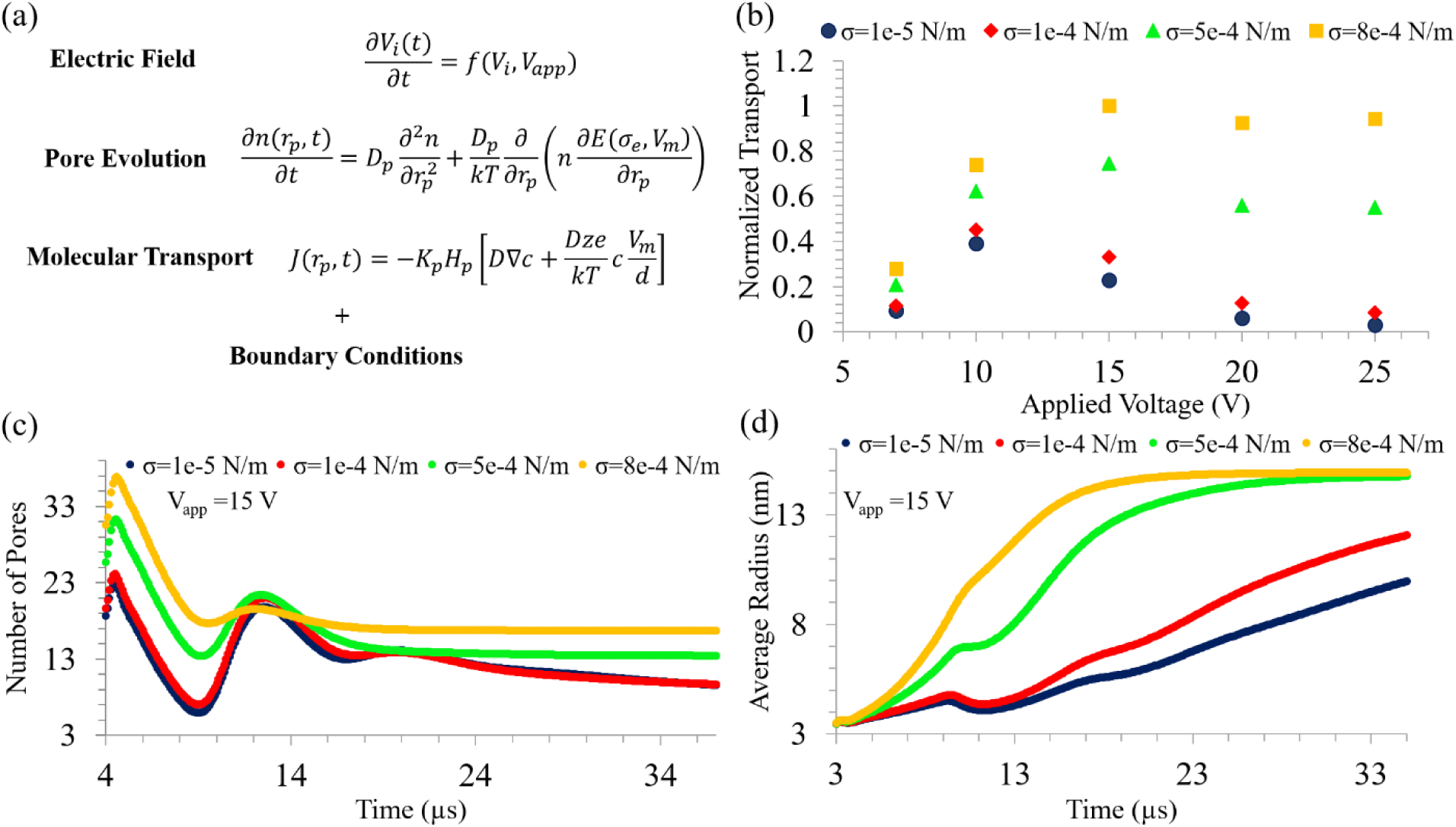
Results obtained from numerical simulations (a) The governing equations describing the Electric Field, Pore Evolution and Molecular Transport. These coupled equations are solved with appropriate initial and boundary conditions to obtain the transported amount in localized electroporation mediated delivery and sampling (see main text for details), (b) The normalized transport (see equation 6) is plotted as a function of the applied far-field voltage (*V_app_*) and membrane tension (*σ*). The transport increases as *σ* is increased. At lower *σ* there is an optimal intermediate *V_app_* for which the transport is maximum. At higher membrane tensions the transport is more uniform over a wider range of *V_app_*, (c) The number of large pores (> 3 nm) formed during the pulsation period is plotted over time for different values of membrane tensions. At higher membrane tension values, the number of large pores formed is increased and they remain open for a longer period, (d) The average radius of large pores (> 3 nm) is plotted over time. The average radius saturates to the largest radius (*r_max_* = 15 *nm*) in the simulations quickly for higher membrane tensions. This suggests that larger pores exist for a longer duration when the membrane tension is elevated.

#### An optimum voltage exists for maximum molecular transport

Our model also suggests that there is an optimum voltage, especially at lower values of membrane tension (*σ*) for which the transport of large molecules is maximized (**see Figure 2b**). In the simulations, at a low far-field voltage (7 V), a population of small pores (~25 pores) expand to a large size (> 3 nm) and then collapse within 15 µs to smaller radii due to a drop in the TMP and effective membrane tension (*σ_e_*). Only a few large pores (<5 pores) remain open over longer durations. On the other hand, at very high far-field voltages (20 V-25 V), several small pores are created that do not expand to larger radii (> 3 nm), thus hindering the transport of large molecules. Less than 10 large pores are observed in these cases. Only at intermediate far-field voltages (10 V-15 V), sufficient number of stable large pores (~15 pores) are created, which enhances the transport of large molecules (**see Figure 3a and 3b**). To experimentally verify this prediction, we systematically varied the far-field electroporation voltage from 10 to 40 V and monitored the transfection efficiency of an mCherry encoding plasmid in HT-1080 cells and MDA-MB 231 cells. The pulse width and number were kept constant at 5 ms and 100, respectively. We observed the transfection efficiency to be the highest for a voltage amplitude of 30 V (**see Figure 4a-b**). At lower voltages of 10 and 20 V, the amount of plasmid delivered was sub-optimal (**see Supplementary Figure 2**), as is evident from the lower fluorescence intensity. For a voltage amplitude of 40 V, in addition to weak fluorescence intensity, the cell viability was low and the morphology appeared abnormal.

**Figure 3:**
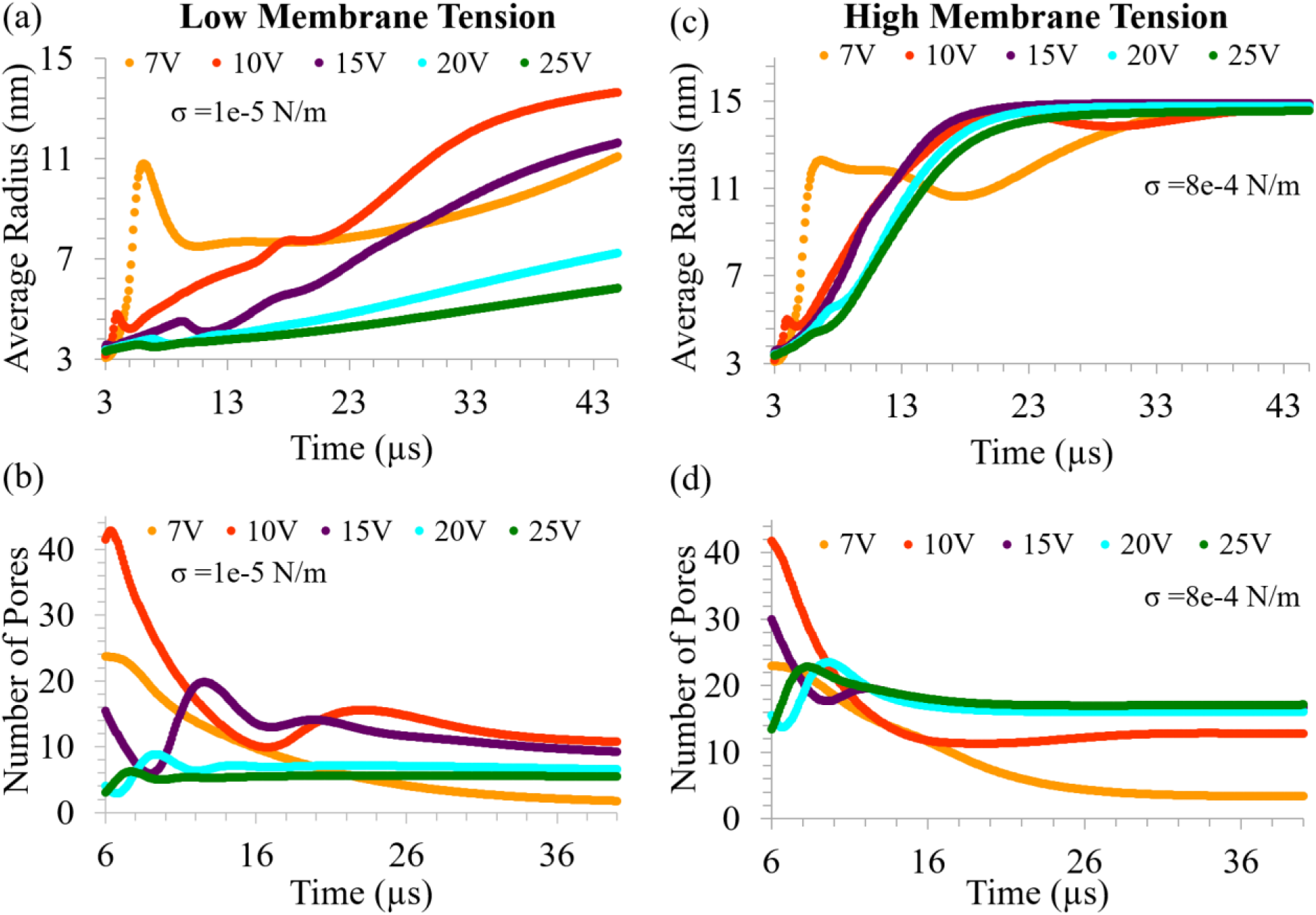
Dependence of pore dynamics on applied voltage (a) The average pore radius of large pores (> 3 nm) is plotted over time for different applied voltages (*V_app_*) and for a low membrane tension value (1e-5 N/m). It is seen that the average radius is large and is sustained over a longer period for an intermediate voltage of 10 V. At a lower voltage (7 V) the average radius is high initially but drops thereafter. For higher voltages the average radius is lower indicating that the pores do not expand to a large size, (b) Corresponding plot of the number of large pores (> 3 nm) formed over time for low membrane tension (1e-5 N/m). At an intermediate voltage of 10 V, many large pores are formed and sustained for a longer duration, (c) The average pore radius of large pores (> 3 nm) is plotted over time for different applied voltages (*V_app_*) for a high membrane tension value (8e-4 N/m). The trend of mean radius is uniform over a range of *V_app_* (10 V-25 V), (d) Corresponding plot of the number of large pores (> 3 nm) formed over time for high membrane tension (8e-4 N/m). Similar number of large pores are formed over a range of *V_app_* (10 V-25 V).

**Figure 4:**
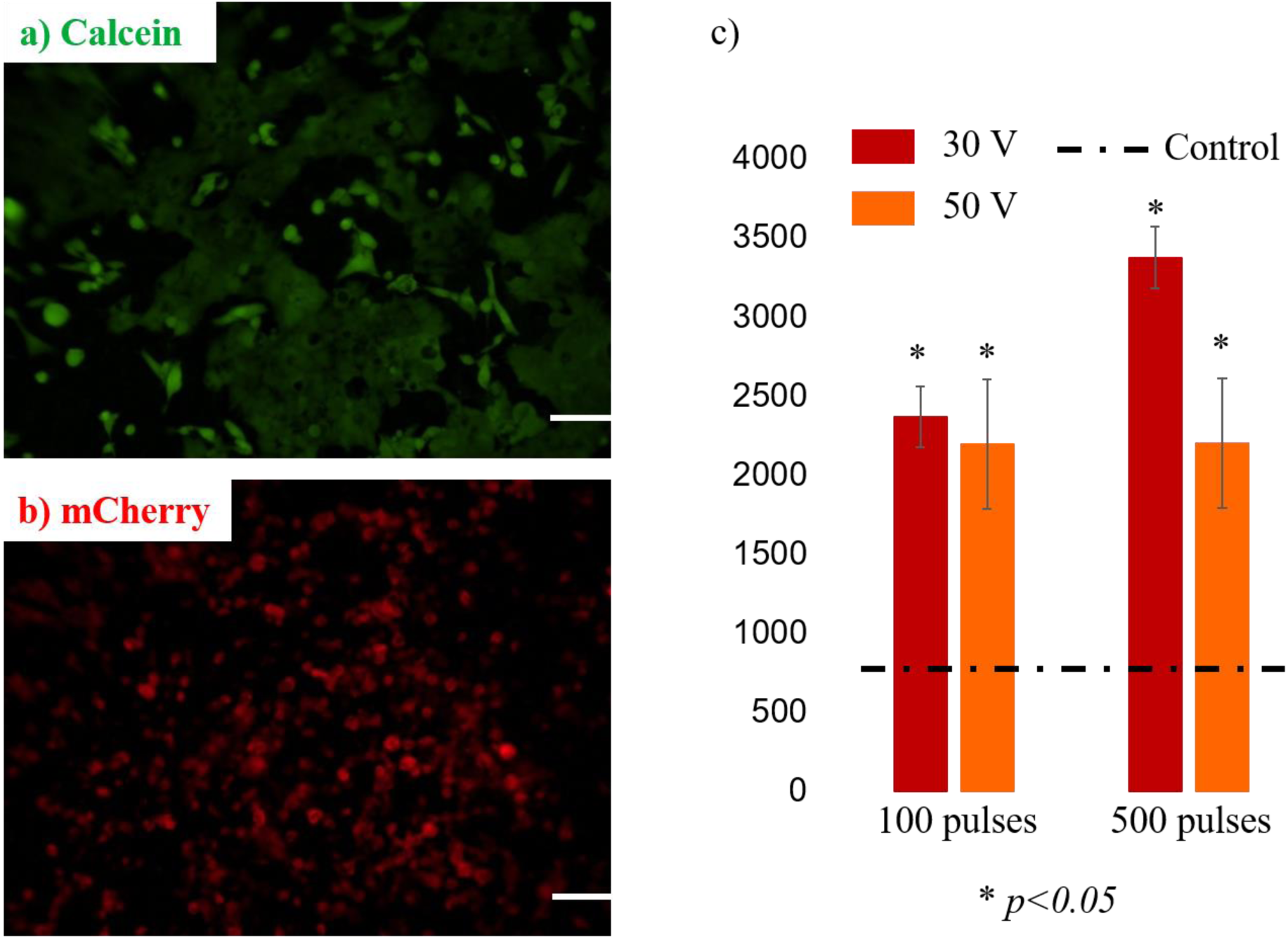
Transfection of mCherry plasmid and tdTomato sampling in MDA-MB 231 cells. a) Calcein AM stained MDA-MB 231 cells 24 hours post electroporation showing high cell-viability. b) Fluorescence image of mCherry plasmid expression in the cells, 24 hours post electroporation. The optimum voltage amplitude of 30 V was used in these experiments. Scale bars = 50 µm. c) Relative Fluorescence Intensity (y axis) of the sampled tdTomato under various electroporation parameters. Media control is shown as a dotted line. The reported error bars are standard deviation (SD) values from n=3 experiments in each case

As a further validation, we investigated how the extracted amount of exogeneous tdTomato protein varied as a function of the voltage amplitude and pulse number in engineered MDA-MB 231 cells. For a voltage amplitude of 30 V, the amount of tdTomato sampled increased with the number of pulses (**see Figure 4c**). On the contrary, when the voltage amplitude was increased to 50 V, the sampling amount was comparable to the 30V, 100 pulses case and independent of the pulse number. The plasmid transfection and tdTomato sampling data taken together highlight the existence of a critical voltage for optimal molecular influx and outflux. We also investigated cell viability on days 1, 2 and 3 post electroporation using live-dead staining. While the cells that were electroporated with a voltage amplitude of 30 V showed healthy morphology, normal cell proliferation and high viability (>99%, **see Figure 5**), most cells electroporated using 50 V had detached from the substrate on day 1 itself. (**see Supplementary Figure 4**). The cells that remained on the substrate showed an unhealthy morphology with blebbing and a spotty tdTomato expression.

**Figure 5:**
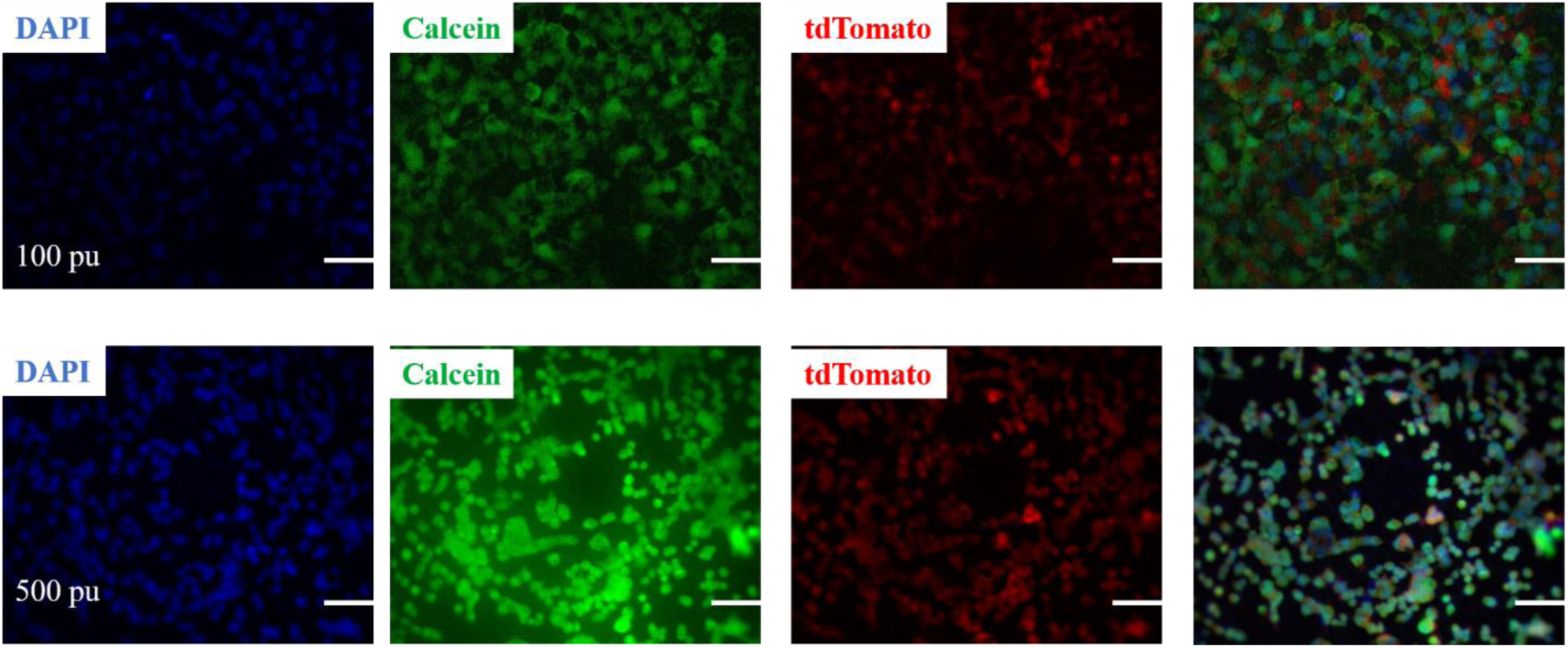
Day 3 viability of cells electroporated with a voltage amplitude of 30 V and 100 pulses (top row), 500 pulses (bottom row). The last image in each row is a composite of the blue, green and red images. Scale bars = 50 µm.

Similar trends have been observed in the case of bulk electroporation. Experimental demonstrations using bulk electroporation systems have shown that very strong pulses are not conducive to large molecule delivery[63]. Subsequent numerical calculations in the context of bulk electroporation have predicted that beyond a certain critical voltage the pores created are small in size[40].

#### Molecular transport is uniform over a broader voltage range for higher cell membrane tension

Interestingly, the model predicts that at higher membrane tensions (*σ*) the molecular transport is uniform over a wider range of applied voltages (**see Figure 2b**). This is a direct result of the fact that at higher membrane tensions the average pore radius and the number of large pores (>3 nm) formed have smaller variability across the applied voltage range (**see Figure 3c and Figure 3d**). To validate this prediction, we delivered Alexa Fluor 488 conjugated BSA (molecular weight = 66.5 kDa, radius ~ 3nm) into tdTomato expressing MDA-MB 231 cells under both hypo-osmolar (~90 mOsmol/kg) and iso-osmolar (~280 mOsmol/kg) buffer conditions. Hypo-osmolar conditions increase the membrane tension by inducing osmotic swelling of the cells [58]. The applied voltage was maintained at 30 V for both cases. We found that under hypo-osmolar conditions the fluorescence intensity of delivered BSA was higher and more uniform as compared to the iso-osmolar case (**see Figure 6a**). By plotting the fluorescence signal from 30 cells that were randomly chosen across three biological replicates, we found the mean signal to be higher and the variability across cells to be lower when the hypo-osmolar buffer was used (**see Figure 6b**). It is worth mentioning that without an accurate measurement of membrane tension it is not possible to directly correlate our experimental results to the model predictions. However, the most pertinent inferences from the model are recapitulated in the experiments. In addition, we observed that cells expressing higher tdTomato fluorescence intensity had lower BSA content and vice versa (**see Figure 6a**). This suggests that the process of delivery and sampling from the cytosolic milieu are directly related. Indeed, effectively permeabilized cells on the nanochannel LEPD substrate uptake BSA through the electro-pores but also lose their cytosolic tdTomato to the surrounding media in the process.

**Figure 6:**
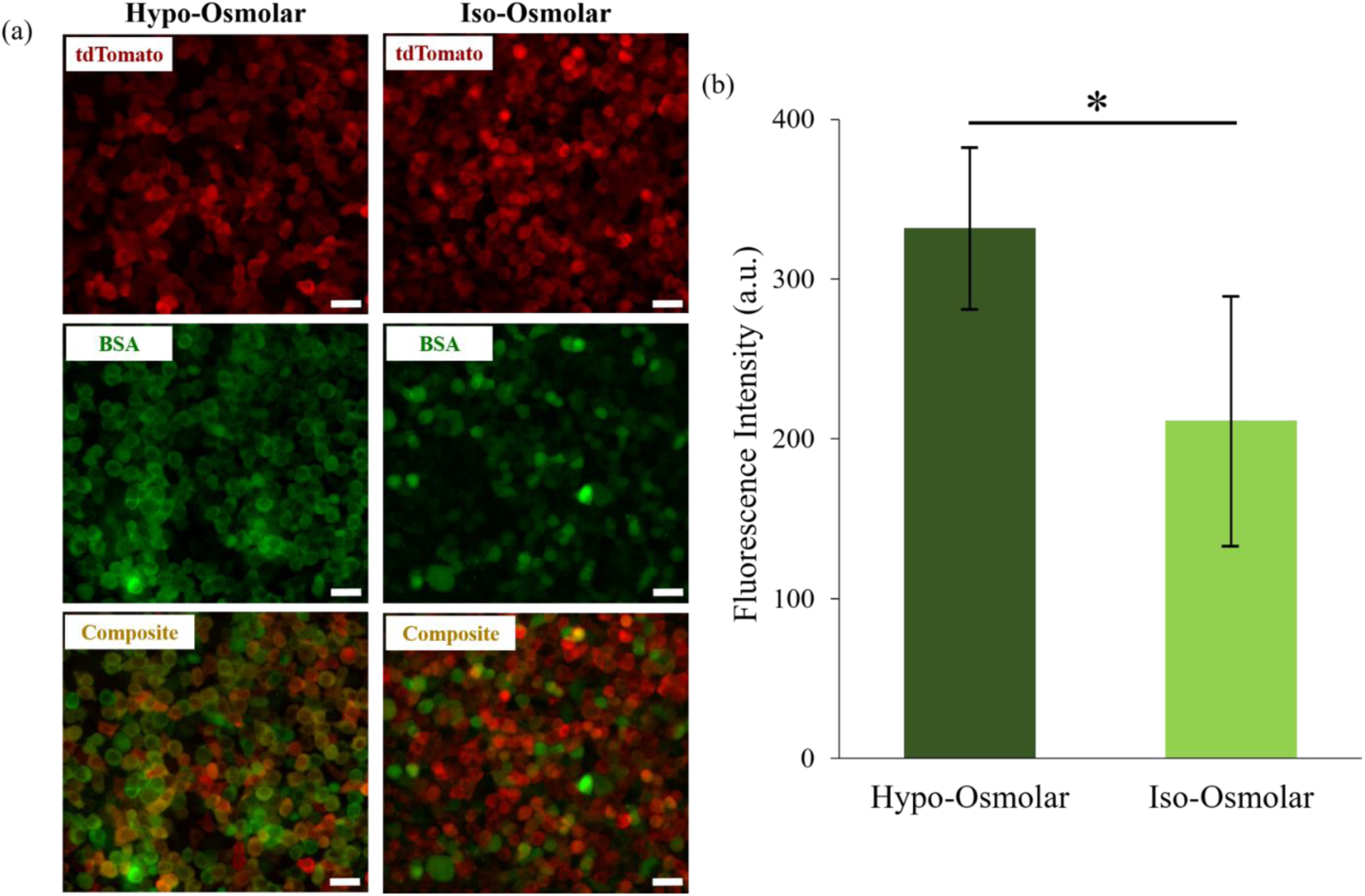
Delivery of BSA into tdTomato expressing MDA-MB 231 cells on the LEPD (a) Comparison of BSA delivery under hypo-osmolar and iso-osmolar conditions. Top fluorescence micrographs (red channel) show TdTomato expression in the cells after electroporation. Middle fluorescence micrographs (green channel) show the efficiency of BSA delivery under hypoosmolar and iso-osmolar conditions. Bottom images are a composite of the red and green channels showing that tdTomato and BSA expressions are inversely related. This indicates that tdTomato has been sampled from cells in which BSA has been efficiently delivered (All scale bars = 50 µm), (b) The fluorescence intensity (in arbitrary units) of BSA is plotted for cells electroporated under hypo-osmolar and iso-osmolar conditions. Error bars indicate the standard deviation (SD). Increased delivery efficiency is observed for the hypo-osmolar case (n=30, *p<0.05).

It is important to note that although the applied far-field voltage across the LEPD is fixed for a particular experiment, factors such as device architecture, cell shape or size and spatial variation in the applied electric field can lead to variability in the transmembrane potential across individual cells. At a higher membrane tension, the molecular transport is less sensitive to the strength of the applied electric field, as a result of which the variability induced by local fluctuations in the transmembrane potential is alleviated. This conclusion is relevant in the context of delivery and sampling via electroporation. This lack of uniformity and efficiency is a major source of technical noise for both bulk and micro/nano electroporation platforms [64], which hinders translation to practical applications such as the study of biological heterogeneity at the single-cell level. However, by optimizing device design and increasing the membrane tension, this variability can be minimized, allowing for improved accuracy and reliability.

#### Small molecule transport is less sensitive to cell membrane tension

We have seen from the model predictions that the delivery or sampling of large molecules may be sub-optimal at lower membrane tensions unless optimal voltage parameters are applied. Increasing the membrane tension increases the amount and uniformity of transport. However, the transport of small molecules (< *r_min_*) is less sensitive to the membrane tension and efficient transport can be achieved even at lower membrane tensions. The transport of small molecules increases linearly with the applied voltage and shows less variation with membrane tension as compared to large molecule transport (**see Supplementary Figure 1**). This was confirmed by our ability to efficiently deliver Propidium Iodide (PI) (hydrodynamic radius ~0.6 nm) into HT 1080 cells without increasing the cell membrane tension, at different applied voltages (**see Figure 1e and Supplementary Figure 1**). Insensitivity to membrane tension and linear increase with applied voltage has also been reported for the delivery of small molecules in the case of bulk electroporation[65]. Another factor contributing to the efficient delivery of small molecules is the presence of pores (<1.5 nm) that are open even after the pulsation period as seen in the model and experimentally verified by Co^2+^ quenching of Calcein AM (**see Supplementary Figure 3**).

## CONCLUSION

In our current work, we have developed a multiphysics model of localized electroporation and molecular transport to explain the delivery and sampling of molecules in live cells. Using this model, we have investigated the role of cell membrane tension and the applied electric field strength in determining the molecular transport. Our model predicts that higher membrane tension increases the amount and uniformity of molecular transport for large molecules over a broad range of applied far-field voltages. Furthermore, we observed that an intermediate voltage is optimal for the transport of large molecules (> 3nm hydrodynamic radius). However, the transport of small molecules is less sensitive to membrane tension and increases linearly with applied voltage. We experimentally validated the model predictions by delivering small molecules such as PI and Co^2+^ as well as a larger protein (BSA) and plasmid (mCherry) into cells using the LEPD system. Finally, we demonstrated the sampling of a large cytosolic protein (TdTomato) in an engineered cell line using optimal parameters for the LEPD without compromising cell-viability. Overall, our simulation and experimental results suggest that localized electroporation in the LEPD is a promising method of non-destructive temporal sampling of cells. Using optimal experimental parameters obtained from the developed model, localized electroporation can be used for *non-destructive, temporal single-cell sampling*, which can provide several advantages over existing techniques. Current single cell profiling techniques enable us to comprehensively identify the molecular state of individual cells. However, these methods are destructive and are unable to track gene expression in the same cells over time. To overcome this issue, temporal dynamics is inferred computational from the single timepoint data. Due to the inherent asynchrony and heterogeneity of a cell population, single-cell data inevitably consist of a large distribution of molecular states. From these data, it is possible to construct mathematical models that build time trajectories of cell fate. These models assume maximum parsimony and as such, the cellular states observed are connected by a trajectory involving minimal transcriptional changes [66]. However, the dynamics of cell state may be non-hierarchical or stochastic [66] and the maximum parsimony principle may not always hold. This effectively means that a single starting point can lead to multiple branching trajectories or multiple trajectories can lead to the same distribution of states, making it possible to describe a molecular state through several regulatory mechanisms[67]. In order to distinguish between alternative dynamical pathways and provide additional constraints that enable the identification of the true underlying mechanisms, anchor points or known cell states in the differential pathways are essential [66]. Emerging methods partially address this issue by combining high-throughput single cell sequencing methods with Cas9 or barcode based genetic perturbations to track cell lineage during organism development [68, 69]. However, to understand the decision-making pathways involved in single-cell processes such as cell differentiation and maturation, tracking the same cell in time is necessary. Temporal sampling of single cells can provide critical information necessary to determine the anchor points involved in these pathways. By combining a high-throughput microfluidics platform for localized electroporation, state of the art ultra-sensitive assays and insights gained from the developed model to optimize experimental conditions, non-destructive sampling and analysis of single cells can be realized in the future.

## ACKNOWLEDGEMENTS

This work was supported by the National Cancer Institute of the National Institutes of Health (NIH) under Award Number U54CA199091 and by NIH SBIR R44 GM110893-02. The content is solely the responsibility of the authors and does not necessarily represent the official views of the National Institutes of Health.

## MATERIALS AND METHODS

### Device Fabrication and Assembly

The device was fabricated using standard soft lithography technique. Briefly, polydimethylsiloxane (PDMS) prepolymer (Sylgard 184, Dow Corning) was mixed in 10:1 (w/w) ratio (base, curing agent), poured on a flat polystyrene dish and cured in an oven at 80°C for 4 hours. The volume of the poured mixture was adjusted to obtain 2 mm thick PDMS slabs. The slabs were cut into 2 cm × 2 cm squares using a razor blade. Holes of the desired diameter were punched in the PDMS slabs using biopsy punches (Accuderm) to form the cell-culture well and delivery/sampling chamber of the device (**see Figure 1a**). A thin layer of uncured PDMS was prepared by spin-coating the mixture at 1500 rpm for 2 minutes on a silicon wafer. The top cell-culture wells and the bottom delivery/sampling chambers were stamped onto the wafer to ink their surfaces with the uncured PDMS. A 13 mm polycarbonate (PC) filter membrane (5×10^8^/cm^2^ porosity, AR Brown) was sandwiched between the two layers and cured in the oven at 80°C for 2 hours to obtain the assembled devices (**see Figure 1b**). The devices were sterilized by incubating in 70% ethanol for 15 minutes, rinsing thoroughly with DI water and drying. This was followed by UV exposure for 1 hour. All subsequent steps were carried out in a laminar flow hood to maintain sterility. The top surface of the PC membrane was coated with Fibronectin (Sigma-Aldrich) by incubating in a 1: 50 (v/v) solution (0.1% Fibronectin, 1×PBS (Gibco™)) to promote cell adhesion. The bottom PC surface and the delivery/sampling chamber were then passivated by dipping the device in a 0.2% (w/v) solution of Pluronic F-127 (Sigma-Aldrich) in 1×PBS. The device was then washed 3 times with 1×PBS to remove any unattached residues before seeding cells.

### Multiphysics Modeling

Numerical simulations were performed in COMSOL Multiphysics 5.2a. A lumped circuit model (**see Figure 1c**) including passive electronic components was used to solve for the transient electric field. The governing coupled ordinary differential equations (ODEs) for the electric field were solved using the Global ODEs and the differential algebraic equations (DAEs) module, available in COMSOL, to obtain the transmembrane potential (TMP). The formation and evolution of pores on the plasma membrane in response to the applied TMP is governed by the Einstein-Smoluchowski equation[70]. A tension coupled non-linear form of this equation[55, 56], which accounts for the tension mediated interaction between pores, was solved using the General Form partial differential equation (PDE) module to obtain the distribution of electro-pores at every time point. The electric field and the pore evolution equations were coupled through the effective electrical conductivity (κ) of the cell membrane. The electro-diffusive transport of molecules[43] across the cell membrane during electroporation was estimated using the Nernst-Planck equation (see **RESULTS AND DISCUSSIONS** and **Figure 2a**). For the Finite Element Analysis, quadratic Lagrange shape functions were used for spatial discretization. The discretized non-linear equation was solved using the Newton-Raphson method at every time step. Temporal integration was performed using the implicit generalized-α scheme.

### Cell Culture

HT1080 and MDA-MB 231 cell lines were obtained from the Mrksich Lab at Northwestern University. CHO and TdTomato expressing MDA-MB 231 cell lines were obtained from Recombinant Protein Production Core (rPPC) and Developmental Therapeutics Core (CDT) facilities at Northwestern University. HT1080, MDA-MB 231 cells were cultured in DMEM (Gibco™) supplemented with 10% FBS (Gibco™) and 1% Penicillin-Streptomycin (Gibco™). CHO cells were cultured in DMEM/F-12 (Gibco™) supplemented with 10% FBS and 1% Penicillin-Streptomycin. The cultures were passaged every 3-5 days upon reaching 80-90% confluency using 0.25% Trypsin (Gibco™). The cells were plated on the device by dispensing 30 µl of cell suspension at a desired density and allowed to adhere. The devices were placed within a 6-well plate (USA Scientific) with the appropriate medium depending on the cell type, inside the incubator (at 37°C with 5% CO2) for a day before performing the electroporation experiments. All experiments were performed on cultures that were passaged less than 10 times.

### Electrical Setup and general protocol for electroporation

A function generator (Agilent) connected to a voltage amplifier (OPA445, Texas Instruments) was used to apply the electroporation pulses (10-50 V, 1-5 ms square pulses, 100-500 pulses, 1-20 Hz). The voltage traces were verified on an oscilloscope (LeCroy). Two ITO coated glass slides (top and bottom, **see Figure 1a**) served as the positive and ground electrodes for pulse application. The molecular cargo for delivery was loaded in the bottom chamber of the LEPD. Conversely, the extracted molecules in sampling experiments were collected from the same chamber and transferred to a 96-well plate.

### Delivery of small molecules (PI and Co^2+^)

MDA-MB 231 cells (~ 10,000) plated in the LEPD on the previous day were first washed three times to remove debris before loading 1xPBS in the cell culture chamber. Propidium Iodide (Life Technologies, diluted to a concentration of 20 µg/ml in 1×PBS) was loaded into the bottom chamber. The device was placed between the two ITO coated glass slides (Nanocs) and three pulses of 0.5 ms duration and 15 V amplitude were typically applied in these experiments. After waiting for 15 mins to allow for molecular diffusion, the LEPD was washed 3 times with 1×PBS to remove residues before imaging on a fluorescence microscope.

For Co^2+^ quenching of Calcein-AM, CHO cells (~10,000) plated on the LEPD the previous day were first stained with Calcein-AM (Life Technologies, diluted to 1 µg/ml in 1×PBS) and incubated for 15 minutes. The cells were washed thrice before loading with 1×PBS. The bottom chamber was also loaded with 1×PBS and electroporation pulses (15 V, 0.5 ms, 1-3 pulses) were applied. 500 mM CoCl_2_ was introduced approximately 60 s post-electroporation to visualize the loss of green fluorescence. The quenched fluorescence was later recovered by injecting 1 mM EDTA solution 3 minutes post electroporation[71].

### Delivery of proteins (BSA)

BSA delivery was performed using a 2.5 mg/ml solution of BSA with Alexa Fluor™ 488 conjugate (Invitrogen) in 1×PBS or 1×Hypo-osmolar buffer.

### Delivery of Plasmids

A 4-kb plasmid encoding the fluorescent protein mCherry (gift from Dr. Vincent Lemaitre, iNfinitesimal LLC) was used at a concentration of 20 ng/µl in 1×Hypo-osmolar buffer. Approximately 10,000 MDA-MB 231 or HT 1080 cells were plated on a device and allowed to adhere for 24 hours before electroporation. The electroporated cells were then washed thrice with 1×PBS and incubated for a day (at 37°C and 5% CO2) before imaging on a fluorescent microscope. An electroporation train of 100 voltage pulses, each of 5 ms duration, was applied at a repetition frequency of 20 Hz. The voltage amplitude was varied from 10-40 V to optimize the efficiency of transfection.

### tdTomato Sampling

tdTomato expressing MDA-MB 231 cells were plated at a density of 10,000 cells per device and incubated for 24 hours. Both the top and bottom chambers in the device were loaded with 1×Hypo-osmolar buffer before applying electroporation pulses. In different experiments, the pulse amplitude and duration were kept constant at 30 V or 50 V and 5 ms respectively, while the number of pulses was varied from 100 to 500. The sampled tdTomato molecules were allowed to diffuse to the bottom chamber for 30 minutes before transferring them to a 96-well plate using a micropipette. A well plate reader (Synergy H1m) was used with the following settings to measure the fluorescence level from the extracted protein – excitation/emission at 553/582 nm, gain: 150 and integration time 1 sec.

### Fluorescence Microscopy and Image Analysis

Fluorescence images were acquired on a Nikon Eclipse Ti-U Microscope equipped with an Andor Zyla 5.5 sCMOS camera. Image acquisition was controlled using Micro-Manager software[72]. The acquired images were analyzed using FIJI, an open source image-processing package[73].

### Viability Analysis

For viability analysis, the cells were stained with Calcein AM (Sigma-Aldrich) and Hoescht 33342 (Life Technologies) using standard protocols. Cells expressing Calcein AM and Hoescht fluorescence simultaneously were alive while the ones expressing only Hoescht were dead.

